# Hypoxia inducible factor HIF1α elevates expression of mRNA capping enzyme during cobalt chloride-induced hypoxia

**DOI:** 10.1101/2024.07.05.602207

**Authors:** Safirul Islam, Chandrama Mukherjee

**Affiliations:** Institute of Health Sciences, Presidency University, Plot No. DG/02/02, Premises No. 14-0358, Action Area 1D, Kolkata 700156, India

**Keywords:** . mRNA capping enzyme/RNGTT, cytoplasmic capping, CoCl_2_ induced hypoxia, HIF1α, long noncoding RNA

## Abstract

In response to hypoxia, hypoxia-inducible factors (HIFs) control the transcriptomic output to mitigate the hypoxic stress. Long noncoding RNAs (lncRNA) are found to be very crucial in regulating hypoxia. Like mRNAs, lncRNAs are protected by 5’ caps that are added by mRNA capping enzyme (CE) in the nucleus. The previous concept that capping takes place in the nucleus was changed by the recognition of a cytoplasmic pool of capping enzyme (cCE). cCE has been shown to recap its substrate uncapped mRNAs or long noncoding RNAs (lncRNAs) present in the cytoplasm, preventing their degradation, even during arsenite-induced oxidative stress. In this study, we examined the effect of CoCl_2_ induced hypoxia on cCE and its function in regulating the substrate lncRNAs. Here, we show that CoCl_2_ induced hypoxia elevates the expressions of nuclear and cytoplasmic CE in HIF1α dependent manner as evidenced by Chromatin immunoprecipitation and HIF1α inhibitor experiments. Furthermore, we found cCE post-transcriptionally controls the stability of its target lncRNAs amidst CoCl_2_ induced hypoxia. These results suggest that cCE, upregulated by HIF1α, may act as a posttranscriptional modulator for a few cCE-targeted lncRNAs.

## 1. Introduction

Hypoxia is one of the contributing factors for generating a solid tumor microenvironment and plays a key role in the progression of tumor metastasis. In 1953 *Gray et. al.* showed the responsiveness of hypoxic tumor cells toward radiotherapy is poor as compared to well-oxygenated tumor cells (1). Additional studies revealed that hypoxia has a profound effect on tumor cell biology which includes its metabolism, angiogenic potential, and sensitivity to chemotherapy or immunotherapy (2). Oxygen-sensitive hypoxia-inducible factor 1α (HIF1α) is the master regulator of hypoxia which reprograms the cellular transcriptomic output in concert with constitutively expressed β subunit so that cells can withstand hypoxic stress (3–5). In addition to HIF1α, several noncoding RNAs like microRNAs (miRNA), long noncoding RNAs (lncRNA), circular RNAs (circRNA)) and epigenetic factors also regulate the cellular response to hypoxia (6).

Cellular stresses are categorized into two groups type 1 and type 2. Hypoxia, heat shock, and arsenite treatment belong to type 1 stress whereas genotoxic drugs belong to type 2 stress. Cells initiate specific cell stressor mechanisms for their survival upon confronting any environmental stress (7). Studies have indicated that hypoxia markedly reduces *de novo* RNA synthesis mediated by modulating chromatin architecture (8,9). In addition, cellular response to hypoxia is also achieved at the post-transcriptional level which includes mRNA splicing, stability, etc. Hypoxia-induced transcriptional regulation is well studied whereas its effect on RNA stability at the post-transcriptional level is still not comprehensive. The 5’ cap is one of the crucial factors that determines the stability of an RNA transcript and how this structure is affected during hypoxic stress, remains elusive.

The majority of eukaryotic RNAs that are transcribed by RNA pol II must have a 5’ cap structure which consists of an N^7^ methylated guanosine residue that is attached covalently to the first nucleotide of the transcripts by an enzyme called RNA guanylyltransferase and 5’-phosphatase (RNGTT) or mRNA capping enzyme (CE) (10,11). Canonical capping takes place inside the nucleus where CE in complex with RNA guanine 7-methyl transferase (RNMT) and RNA guanine 7-methyl transferase activating subunit (RAMAC) act on the 5’end of the nascent RNA transcripts to add a cap (12–14). Later, a pool of cytosolic capping enzymes (cCE) was identified which can recap the uncapped transcripts in the cytoplasm resulting in their stabilization (15,16). cCE forms a complex with the cytoplasmic pool of RNMT, RAMAC along with an unknown 5’RNA kinase and an adaptor protein Nck1 that provides a platform for the assembly of these components to make active cCE that regulates the stability and translability of certain mRNAs and thus has been speculated in maintaining ‘cap homeostasis’ to regulate gene expression (16–18). Inhibiting cytoplasmic capping by overexpressing a dominant negative mutant of cCE (K294A) increased a pool of uncapped RNAs (16,19). In agreement with these findings, studies across several labs revealed the presence of uncapped RNAs in various cells pointing to the possibilities of cytoplasmic recapping. Indeed, with the advancement of imaging techniques, mRNAs have been shown to undergo cyclical decapping and recapping (20). Later, recapping was revealed to act as a nexus of post-transcriptional control as cCE not only regulates mRNAs but also a subset of lncRNAs (21).

Besides these canonical functions of CE/cCE either in the protection and maturation of RNA pol II transcribed transcripts or recapping cytosolic transcripts, recent studies have indicated its potential role in various biological processes ranging from modulating the Hedgehog signaling pathway during *Drosophila* development to its involvement in certain cancers (22,23). A current study demonstrates CE governs biliary-derived liver regeneration initiation by transcriptional regulation of mTORC1 and Dnmt1 in Zebrafish (24). Additionally, one recent study uncovers that METTL3 regulates the expression and activity of CE in non-small cell lung cancer (NSCLC) cell lines (25). Recently, a study revealed the role of cCE in protecting the cap status of its target mRNAs during arsenite-induced oxidative stress (26). However, there is no study regarding the effect of hypoxic stress on CE/cCE.

In this study, we examine the effect of Cobalt chloride (CoCl_2_) induced hypoxia on the expression of CE or cCE for the first time. CoCl_2_ is a well-known hypoxia-mimetic chemical compound that triggers hypoxia-like conditions in vitro (27). Using chemically induced hypoxia on osteosarcoma cell lines (U2OS and MG63) as our model systems, we found CoCl_2_ induced hypoxia elevates the expression of CE/cCE. Furthermore, we examine the steady state level of previously validated cCE-targeted lncRNAs (21) in CoCl_2_ induced hypoxic conditions in cells expressing a dominant negative mutant of cCE (K294A) (21). Our study shows some of these lncRNA are upregulated in CoCl_2_ induced hypoxia and cCE regulates their stability indicating its role in post-transcriptional regulation of a subset of hypoxia-responsive lncRNAs.

## 2. Materials and methods

### 2.1 Cloning of K294A in pLVX-TetOne-Puro vector

Mouse cytoplasmic restricted catalytically inactive capping enzyme (K294A) was amplified from pcDNA4/TO-bio-myc-NES-mCEΔNLS (K294A) construct (21) using Q5^®^ High-Fidelity DNA Polymerase (NEB, cat no# M0491L). The pLVX-TetOne-Puro-myc-NES-mCEΔNLS(K294A) construct was prepared by subcloning myc-NES-mCEΔNLS(K294A) into an empty pLVX-TetOne-Puro vector (Takara Bio, cat. No# 631847) using In-Fusion HD Cloning Kit (Takara Bio, cat no# 638909). The construct was further confirmed by Sanger sequencing and is available upon request. The sequences of the cloning primers are listed in Table S1.

### 2.2 Cell culture, treatments, and generation of stable cell lines

Human osteosarcoma cell line U2OS was purchased from the American Type Culture Collection (ATCC), and MG63 from the National Centre for Cell Science (NCCS, Pune). Both the cell lines were routinely cultured in 1X DMEM (Gibco, cat no# 11965092) supplemented with 10% fetal bovine serum (Himedia, cat no# RM1112) and 1% penicillin-streptomycin (Gibco, cat no# 15140-122) in the presence of 5% CO_2_ at 37 °C in a humidified incubator (Eppendorf). Mycoplasma contamination was tested to confirm contamination-free cell culture using a PCR-detection kit (Sigma-Aldrich, cat no# MP0035).

For hypoxia induction *in vitro*, cells were treated with different concentrations (100 µM, 200 µM, and 300 µM) of CoCl_2_ (Sigma Aldrich, cat no# C8661) solution that was made by dissolving in DPBS (Gibco, cat no# 14040182). After 24 hours of treatment, cells were harvested for RNA/protein extraction (28).

HIF1α inhibitor PX478 was procured from the MedChemExpress (cat #HY-10231). 10mM stock of PX478 was prepared by dissolving it in deionized water. Cells were treated with PX478 to a final concentration of 40µM to inhibit HIF1α activity (29).

To generate inducible stable cell lines expressing bio-myc K294A, U2OS cells were transfected with pLVX-TetOne-Puro bio-myc K294A. After 48 hours of transfection, Puromycin (50µg/ml, Sigma Aldrich, cat# P9620) was added to multiple dilutions of transfected cells and transfectants were incubated for clonal selection of the stable cells according to the manufacturer’s protocols till the isolation of the single colonies. After growing a few selected colonies, Doxycyclin (Dox) (1ug/ml) (Sigma Aldrich, cat no# D5207) was added to the selected clones for inducible expression of K294A. The stable cell line expressing the highest bio myc K294A protein (Fig: S3C-D) was selected for subsequent experiments.

### 2.3 XTT cell viability assay

The viability of osteosarcoma cells upon treatment with different concentrations of CoCl_2_ solution was assessed by using an XTT cell viability assay kit (Cell Signalling Technology, cat no#9095) following the manufacturer’s protocol. Briefly, 2000 cells of U2OS/MG63 were seeded in 96 well plates and treated with different concentrations of CoCl_2_ solutions. 24 hours post-treatment, 50 µl of XTT reagent was added to each well and incubated for 3 h in 5% CO_2_ at 37 °C. The absorbance was measured at 450 nm using a BioTek Synergy microplate reader.

### 2.4 RNA interference and treatment

For Xrn1 knockdown, K294A stable cells uninduced or induced were transfected with Xrn1 siRNA (Ambion, assay ID# s29016) targeting the coding region of *XRN1* and scramble siRNA at 10nM final concentration using Lipofectamine RNAiMAX reagent following manufacturers protocol (Invitrogen, cat no# 13778-075). 24 hrs post-transfection, transfection mix containing media was replenished with fresh growth medium. Total of 48 hrs post-transfection half of the cultured cells were treated with Dox (1µg/ml) to induce K294A expression and maintained in culture for another 24 hrs. Cells were harvested for protein and RNA isolation. The knockdown efficiency was affirmed by western blotting using rabbit anti-Xrn1 antibody (Invitrogen, cat no# PA5-57110).

### 2.5 Cytoplasmic and nuclear fractionations

To isolate cytoplasmic fractions, U2OS cell pellets were resuspended in cytoplasmic lysis buffer (20 mM Tris-Cl of pH 7.5, 10 mM NaCl, 10 mM MgCl_2_, 10 mM KCl, 0.2% NP40, 1mM PMSF, RNAseOUT and 1X protease inhibitor cocktail) and incubated on ice for 10 min with intermittent gentle agitation. Nuclei were removed by centrifugation at 1,000 xg for 10 minutes at 4°C.

To isolate nuclear fractions, the nuclear pellet was resuspended in nuclear lysis buffer (150 mM NaCl, 1% Triton X-100, 0.5% Sodium Deoxycholate, 1% SDS, and 25 mM Tris-Cl of pH 7.5) for 40 minutes with occasional agitation. The nuclear extract was prepared by centrifugation at 10,000 xg for 10 minutes at 4 °C.

### 2.6 RNA extraction and poly(A) selection

Total RNA was extracted by adding TRIzol to the harvested cells (Invitrogen cat# 15596026) according to the manufacturer’s protocol. To isolate cytoplasmic RNA TRIzol reagent was added to cytoplasmic lysate (2.5) following by manufacturer’s protocol. RNAs were then treated with DNase I (Thermo Scientific, cat no# EN0521) and extracted using phenol: chloroform: isoamyl alcohol (Sisco Research Laboratories Pvt Ltd, cat no# 69031). 2µg of DNA-free RNA was used for poly(A) selection using Dynabeads mRNA DIRECT Kit (Invitrogen, cat no# 61011) according to the manufacturer’s instruction.

### 2.7 Quantitative real-time PCR analysis

2 µg of either total RNA or cytoplasmic RNA was reverse transcribed using Super Script III First Strand cDNA Synthesis kit (Invitrogen cat# 18080051) according to the manufacturer’s instruction. A 1:1 cDNA dilution was done to perform real-time PCR using SsoAdvanced Universal SYBR Green Supermix (Bio-Rad cat no#172-5271) in CFX connect Real-Time instrument (Bio-Rad). The PCR cycle was carried out using the following thermal conditions initial denaturation at 95 °C for 30 s followed by 40 cycles of 95 °C for 10 s and 60 °C for 15 s and a melting curve was, generated by steadily increasing the temperature from 65 °C to 95 °C in 0.5 °C increments with a duration of 2 s/ step. *Rplpo* was used as an internal control. The relative fold change of gene expression was calculated using the comparative CT method (ΔΔCt). Unless specified otherwise, at least three different biological replicates were used. The complete list of primers is listed in Table S1.

### 2.8 Western blotting analysis

Cells were harvested in ice-cold DPBS and the cell pellet was collected by centrifuging at 1000 g for 5 min at 4 °C. For Whole cell lysis, harvested cells were resuspended in whole cell lysis buffer (50 mM Tris-Cl pH 8.0, 150 mM NaCl, and 1% NP-40) supplemented with 1mM PMSF and protease cocktail inhibitor (Roche, cat no# 4693159001) and kept in ice for 30 minutes with intermittent vortex. Following incubation, the whole cell lysate was collected by centrifuging the suspension at 12000 xg for 10 minutes at 4 °C. A Rapid Gold BCA Protein Assay kit (Pierce, cat no# A53226) was used to quantify protein samples. 30-60 µg protein extracts were fractionated using 8-10% SDS-PAGE under reducing conditions and transferred onto Immobilon-FL PVDF (Millipore, cat no# IPFL00010) membranes. The membranes were blocked in 3% skimmed milk in phosphate-buffered saline (PBS) for 30 min at room temperature followed by incubation in respective primary antibody dilution. The dilutions are as follows 1:1000 rabbit anti-HIF1α (Cell Signaling Technology, cat no# 36169); 1:1000 mouse anti-beta actin (Santa Cruz Biotechnology, cat no# SC-517582), 1:1000 rabbit anti-CE (Novus Biologicals, cat no# NPB1-49973); 1:2000 mouse anti-GAPDH (Novus Biologicals, cat#2D4A7); 1:2000 rabbit anti-CA9 (Cloud-clone Crop, cat# PAD076Hu01); 1:2000 mouse anti-lamin (DHSB, cat no# MANLAC1(4A7), 1:1000 rabbit anti-Xrn1 antibodyat 4 °C overnight with constant shaking. Membranes were washed thrice in PBS-T after primary antibody incubation and incubated with secondary antibodies like donkey anti-MouseDyLight 800 (Cell Signaling Technology, cat no#5257) and donkey anti-Rabbit680 (Invitrogen, cat no#A10043) at 1: 10000 dilutions in PBS-T. Blots were scanned in OdysseyCLx Imaging System (LiCor Inc.) and band intensities were calculated using ImageJ software (NIH, USA).

### 2.9 Immunofluorescence

Cells were cultured on coverslips and fixed in freshly prepared fixing solution [4%PFA(EMS)+0.2% Triton-100 in 1X PBS] for 10 min at room temperature. Coverslips were washed thrice in washing solution (0.5mM MgCl_2_, 0.05% Triton-100 in 1X PBS) and blocked in blocking buffer (5% FBS, 0.2M glycine, 0.1% Triton-100 in 1X PBS) for 30 min at room temperature. Coverslips were incubated with primary antibodies (1:100, rabbit anti-HIF1α; 1:100 rabbit anti-CE) overnight at 4 °C. Following washing coverslips were incubated with secondary antibodies of donkey IgG conjugated with either Alexa Fluor 488 (Invitrogen cat no# A21206) or 568 (Invitrogen, cat no# A10037) at a dilution of 1:1000 and Hoechst 33358 (Sigma, cat no# 94403) diluted in blocking buffer for 1 hr at room temperature. Coverslips were mounted on glass slides with a mounting medium (Invitrogen, cat no# S36967). Images were acquired using a Confocal laser scanning microscope (Leica) fitted with a 63X Plan Apo oil immersion objective (NA 1.4). Images were analyzed, and the fluorescence intensity measurements were performed using the Zeiss Zen 3.3 software package.

### 2.10 Chromatin Immunoprecipitation (ChIP) assay

HIF1α binding sites on CE promoter were predicted by using JASPAR software (http://www.jaspar.genereg.net). The ChIP assay was performed using a Simple Chip Enzymatic Chromatin IP Kit (Cell Signaling Technology, cat# 9003) as per the manufacturer’s instructions. Briefly, the treated/ untreated cells were fixed in 1% formaldehyde for 10 min at room temperature to cross-link the DNA-protein complex. Glycine was added to the cell to quench the unreacted formaldehyde. Following washing with ice-cold PBS containing 1mM PMSF, cells were harvested by centrifugation at 2000 g for 5 min at 4 °C. Cell pellet was lysed in lysis buffer (1× buffer A, 0.5 mM DTT, 1× Protease Inhibitor Cocktail, 1 mM PMSF) and nuclei were pellet down at 2000 g for 5 min at 4 °C. The nuclei pellet was resuspended in buffer B supplemented with 0.5 mM DTT and treated with micrococcal nuclease for 20 min at 37 °C. Following centrifugation at 16000 g for 1 min at 4 °C. Nuclei pellet was resuspended in ChIP buffer (1× ChIP buffer, 1× Protease Inhibitor Cocktail, 1 mM PMSF), followed by sonication to break the nuclei membrane and lysate was clarified by centrifugation at 9400 g for 10 min at 4 °C. 10 μg of chromatin DNA was incubated with rabbit anti-HIF1α (Cell Signaling Technology, cat no# 36169) or normal rabbit IgG (Cell Signaling Technology) overnight at 4°C.To the IP samples protein G magnetic beads were added and incubated 2 hrs at 4°C with rotation. The bead-chromatin-antibody complexes were washed three times in a low-salt wash buffer and one time in a high-salt buffer and eluted in elution buffer heating at 65 °C for 30 min. The proteins were digested by treating them with proteinase K at 65 °C for 2 hrs followed by DNA isolation using a spin column. The immunoprecipitated and input DNA were analyzed by Agarose gel electrophoresis as well as subjected to real-time PCR using SsoAdvanced Universal SYBR Green Supermix (Bio-Rad cat#172-5271) in CFX connect Real-Time instrument (Bio-Rad). The primers for ChIP qPCR are listed in Table S1. Relative fold enrichment was calculated using the comparative CT method (ΔΔCt) described previously where IgG was used as the negative control.

### 2.11 *In Vitro* Xrn1 susceptibility assay

30 ng poly(A) selected cytoplasmic RNAs of different experimental conditions from three independent biological replicates were heat denatured at 65 °C for 5 minutes and set up for digestion by adding 0.5 units of Xrn1 enzyme (NEB, cat no# M0338S). Prior to digestion, the cytoplasmic poly(A)RNAs were spiked with synthetic capped Renilla luciferase mRNA as negative control for the experiments respectively. The RNA samples were treated with Xrn1 incubated for 1 hr at 37 °C. After incubation, the digested RNAs were recovered using phenol: chloroform extraction followed by ethanol precipitation. cDNA was prepared using iscript cDNA synthesis kit (Bio-Rad cat no# 1708890). RT-qPCR was performed using gene-specific primer pairs against the 5’-ends of the candidate transcripts as done in earlier studies. The C_t_ values for each cCE targeted transcripts and non-targeted controls were normalized against the C_t_ value of capped Renilla luciferase obtained from each condition. The loss of 5’ ends from each experimental condition was obtained by calculating the differences in normalized C_t_ values of each transcript before and after Xrn1 digestion. ΔX or change in Xrn1 susceptibility in cells was measured by ΔX_K294A_/ΔX_Control_ ratio, where ΔX_Control_ is the relative loss of transcript 5’ ends in control, and ΔX_K294A_ is their relative loss in K294A-expressing cells. The plot showed the relative change in Xrn1 susceptibility between CoCl_2_ treated and untreated cells.

### 2.12 Statistical analysis

The statistical analysis was calculated by performing Two-tail Student’s *t*-tests for two groups and one-way ANOVA for multiple groups using GraphPad Prism 9 Software (GraphPad Software, Inc). All the data were represented as ±SD of at least three independent biological replicates (n=3). Data with p<0.05 were considered statistically significant.

## 3. Result and Discussion

### 3.1 Hypoxia mimetic CoCl_2_ augments HIF1α expression in U2OS and MG63 osteosarcoma cells

The characterization and activity of cCE have been well elucidated in U2OS osteosarcoma cells (15–17,21). Other cell lines like NIH3T3 mouse fibroblasts or human RPE1 (diploid) or primary cells like mouse astrocytes or cardiomyocytes express variable amounts of cCE (26). To determine the effect of hypoxia on the CE/cCE, an *in vitro* chemically induced hypoxic cellular model was established using two osteosarcoma cell lines U2OS and MG63 along with the diploid cell line hTERT RPE-1 (Fig.1, S1). CoCl_2_ is one of the most commonly used hypoxia mimetic reagents for the induction of chemical hypoxia in cellular models and the presence of HIF1α in the cellular extract is the indicator of successful hypoxia induction (28,30,31). Both U2OS and MG63 cells were exposed to different concentrations (100 µM, 200 µM, and 300 µM) of CoCl_2_ solution for 24 h and the induction of CoCl_2_ triggered hypoxia was confirmed by determining the presence of hypoxia marker HIF1α at the protein level by immunoblotting and immunofluorescence staining (Fig. 1). The gradual increase of HIF1α protein expression with the increasing dose of CoCl_2_ at 24 h was observed in both cell lines (Figs. 1A,1B, 1D, & 1E) which affirms the establishment of hypoxia. The toxicity of CoCl_2_ in the cell lines was examined by performing a XTT cell viability assay (32). Concentration-dependent cytotoxicity of CoCl_2_ was observed in both cell lines where 300 µM showed the highest toxicity (Figs. 1C and 1F). Treatment of cells with 200 µM CoCl_2_ concentration did not show any toxicity to both the cell lines. So, considering the expression of hypoxic marker HIF1α and the minimal cytotoxicity of CoCl_2_, 200 µM was selected for further experiments with both cell lines. The immunofluorescence images of HIF1α also corroborate with the immunoblotting results for CoCl_2_ induced hypoxia in U2OS and MG63 cell lines (Figs. 1G and 1H). RPE-1 cells also showed elevated expression of HIF-1α after being treated with 200 µM CoCl_2_ for 24 h (Figs. S1A and B) suggesting a similar mechanism exists.

**Figure 1:**
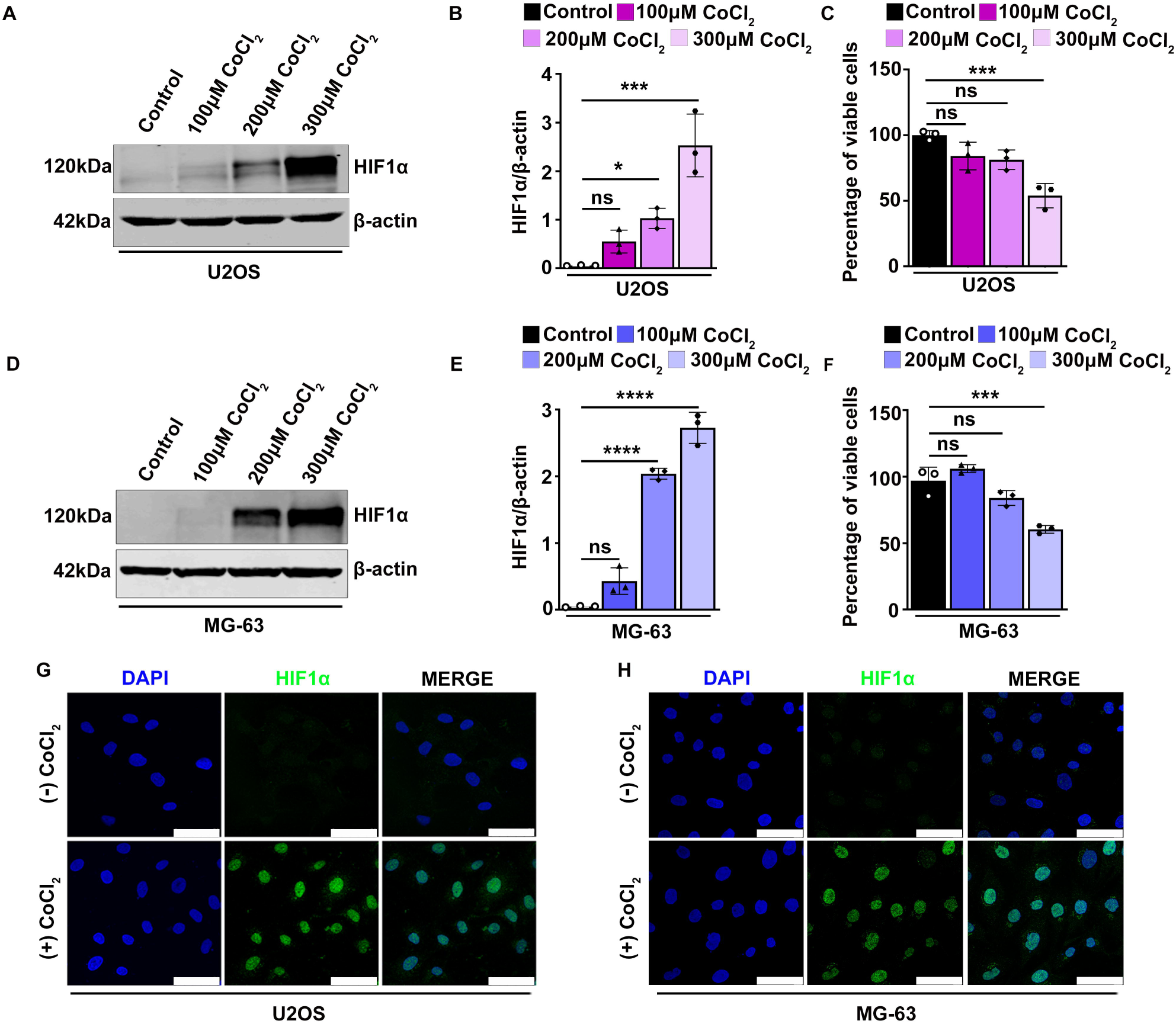
Hypoxia mimetic CoCl. _2_ **augments HIF1α expression in U2OS and MG63 osteosarcoma cells.** A & D. Western blot analysis of hypoxic stress marker HIF1α, showing gradual increase upon increasing concentrations of CoCl_2_ for 24 hrs treatment in U2OS and MG63 osteosarcoma cells respectively. B & E. Quantification of HIF1α protein by ImageJ software using three independent biological replicates. β-actin was used as the loading control. C & F. XTT cell viability assay showing CoCl_2_ dose-dependent loss of cellular viability in U2OS and MG63 cells. G & H. Immunostaining analysis showing an elevated expression of HIF1α (green) during hypoxia in osteosarcoma cells. DNA is represented by DAPI. The scale bar represents 50 µm. Statistical significance was calculated by unpaired two-tailed Student’s t-test represented as ± SD from three biological replicates. *ns, P*: non-significant, *\*P*< 0.05, \*\**P*< 0.005, \*\*\**P*< 0.0005, \*\*\*\**P*< 0.0001; n≥3.

### 3.2 CoCl_2_-induced hypoxia elevates mRNA-capping enzyme expression in osteosarcoma cells

CE protects mRNAs and lncRNAs by adding a 5’ cap structure and besides this protection of 5’ ends of these RNAs it also helps the RNA transcripts for their efficient export from the nucleus to the cytoplasm, co-transcriptional processing, and decay (33,34). However, there are a few studies that report the regulation of expression of mammalian CE itself or its recruitment to target genes. A recent study shows catalytically active CE can bind to RNA polymerase II and its target genes in the presence of c-Myc (35). Earlier, we have shown that CE can shuttle between the nucleus and the cytoplasm and localize to stress granules during arsenite-induced stress (26). However, the impact of other stressors like hypoxia, a major contributing factor to the development of solid tumors, on CE is yet to be known. This led us to examine the effect of CoCl_2_ mimetic hypoxia on CE. To understand this, both U2OS and MG63 cells were incubated with CoCl_2_ for 24 h, followed by staining using CE antibody, and images were captured by a confocal microscope. The immunofluorescence results showed the enhanced expression of total CE, particularly cytoplasmic CE as evidenced by increased staining intensity in the cytoplasm in CoCl_2_ treated cells, compared to untreated cells in both cell lines (Figs. 2A & 2C). We further quantified the mean intensity of CE and our data revealed significant upregulation of CE in CoCl_2_ induced hypoxic cells compared to untreated cells (Figs.2B & 2D). To determine if this increased level of CE was due to the change in the level of the total protein or change in the cytoplasmic-nuclear distribution, total protein levels were measured from untreated and CoCl_2_ treated cells and an increase in the level of CE was observed in both cell lines (Figs: 4B and 4D, discussed later) supporting the immunofluorescence data (Figs. 2A-D). To examine the distribution of nuclear and cytoplasmic CE upon CoCl_2_ treatment, U2OS cells were fractionated in nuclear and cytoplasmic fractions using a standard method (21) and the efficiency of separations were measured by western hybridization with antibodies against Lamin A/C (nuclear protein) and GAPDH (cytoplasmic protein) (Fig: 2E). The chemical induced hypoxic condition was confirmed by the cytosolic expression of CA9, in CoCl_2_ treated cells since it was a positive marker for CoCl_2_ induced hypoxia (36) (Fig: 2E). An increased level of CE was observed in both nuclear and cytoplasmic fractions after CoCl_2_ treatment (Fig: 2E). However, in agreement with the previous immunofluorescence results (Figs. 2A, 2C), cytosolic CE was expressed more in CoCl_2_ induced hypoxia as evidenced by the quantitation (Fig: 2F). A similar increase in the expression of the total population of CE is also observed in RPE1 cells (Fig. S1C-D). Collectively, these data suggest CoCl_2_ induced hypoxia increased expression of CE/cCE in osteosarcoma and human diploid cell lines indicating possible regulation of CE by HIF1α.

**Figure 2:**
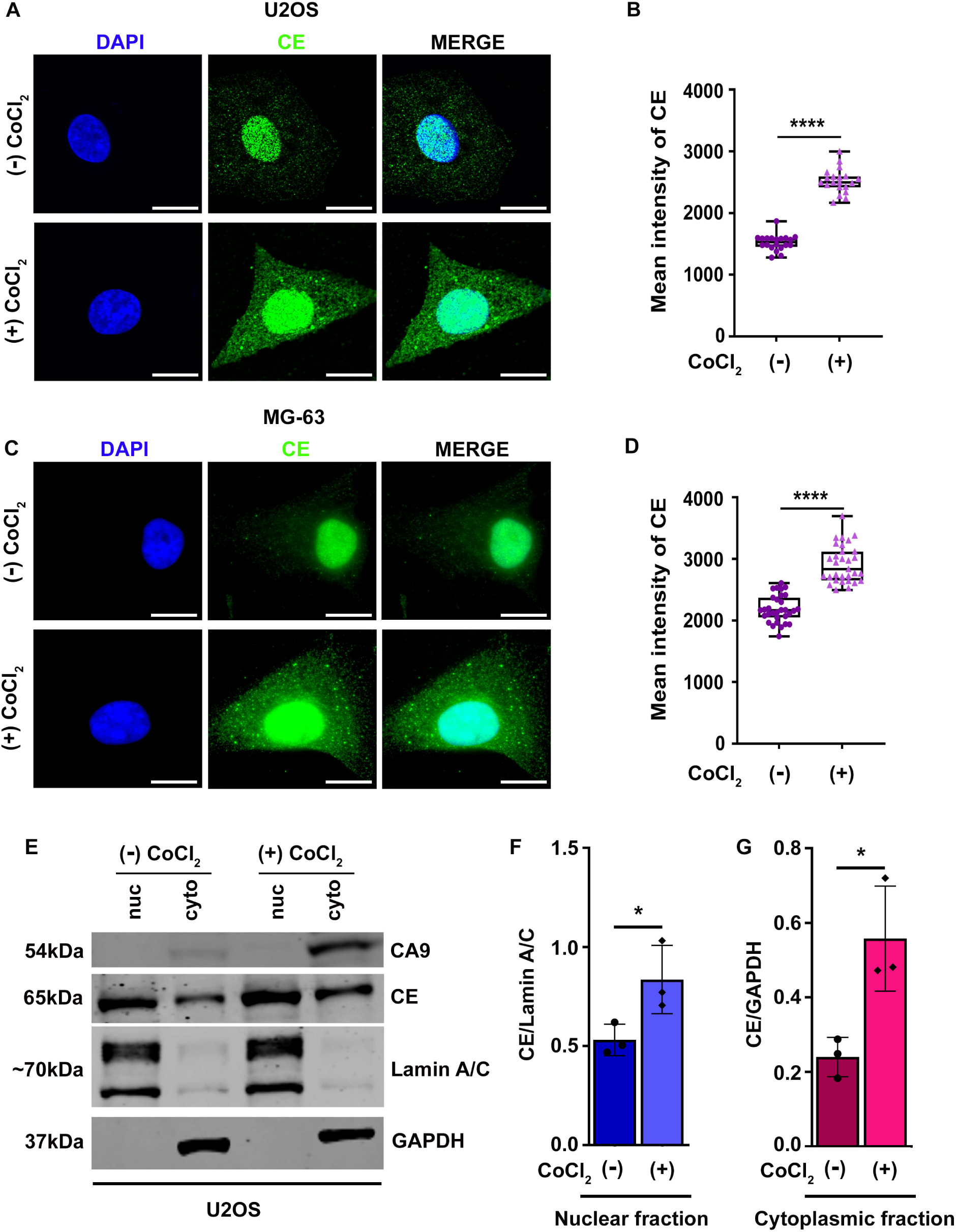
CoCl_2_-induced hypoxia elevates expression of CE/cCE in osteosarcoma cells. A & C. Immunostaining analysis showing enhanced expression of nuclear (CE) and cytoplasmic mRNA capping enzyme (cCE) in CoCl_2_ treated cells as compared to untreated cells. The scale bar represents 50 µm. B&D. Quantification of mean CE intensity using Zeiss Zen 3.3 software package, n≥20 cells. E. Western blot analysis of CE in cytoplasmic and nuclear fractions of U2OS cells treated or untreated with CoCl_2._ GAPDH and Lamin A/C were used as cytoplasmic and nuclear markers respectively whereas CA9 was used as a hypoxic marker. F. Quantification of CE bands intensity from E showing the amount of CE in the nucleus and cytoplasm in U2OS cells treated or untreated with CoCl_2._ Normalization of cytoplasmic and nuclear CE was done using GAPDH and Lamin A/C respectively. Statistical significance was calculated by unpaired two-tailed Student’s t-test represented as ± SD from three biological replicates. *ns, P*: non-significant, *\*P*< 0.05, \*\**P*< 0.005, \*\*\**P*< 0.0005, \*\*\*\**P*< 0.0001; n≥3.

### 3.3 Mammalian *RNGTT* promoter harbours hypoxia-responsive elements (HRE)

Previous studies showed hypoxia or CoCl_2_ induced hypoxia stimulates the expression of HIF1α (37). Similarly, in our CoCl_2_ induced hypoxia model, we observed an accumulation of HIF1α protein during hypoxia (Fig: 1). HIF1α is a member of basic-helix–loop–helix (bHLH)-PAS domain-containing protein family (38) and known to induce the transcription of gene expression by binding to the HRE (R/CGTG) elements present upstream of the transcription start sites in the promoter region (39,40). This led us to speculate that HIF1α might control *RNGTT* expression. *In silico* promoter analysis revealed putative HRE sequences in the promoter of *RNGTT* as represented schematically in Fig. 3A. The *in-silico* data was validated in CoCl_2_ treated or untreated U2OS cells by performing ChIP assay with anti-HIF1α antibody, followed by PCR analysis with different sets of primers targeted to two putative HIF1α binding sites (BS1 & BS2) within 5 kb upstream of the transcription start sites. PCR results showed the direct binding of HIF1α, but not the IgG control to the *RNGTT* promoter in both the binding sites in CoCl_2_ treated U2OS cells but not in untreated cells (Fig. 3B). The absence of binding of HIF1α in the untreated cells may be due to the degradation of HIF1α in the untreated cells as it is unstable in presence of oxygen (41). Further qPCR analysis revealed that in CoCl_2_ induced hypoxic U2OS cells, HIF1α binds to *RNGTT* promoter to BS1 and BS2 with 12-fold and 15-fold enrichment respectively, as compared to untreated cells (Fig. 3C).

**Figure 3:**
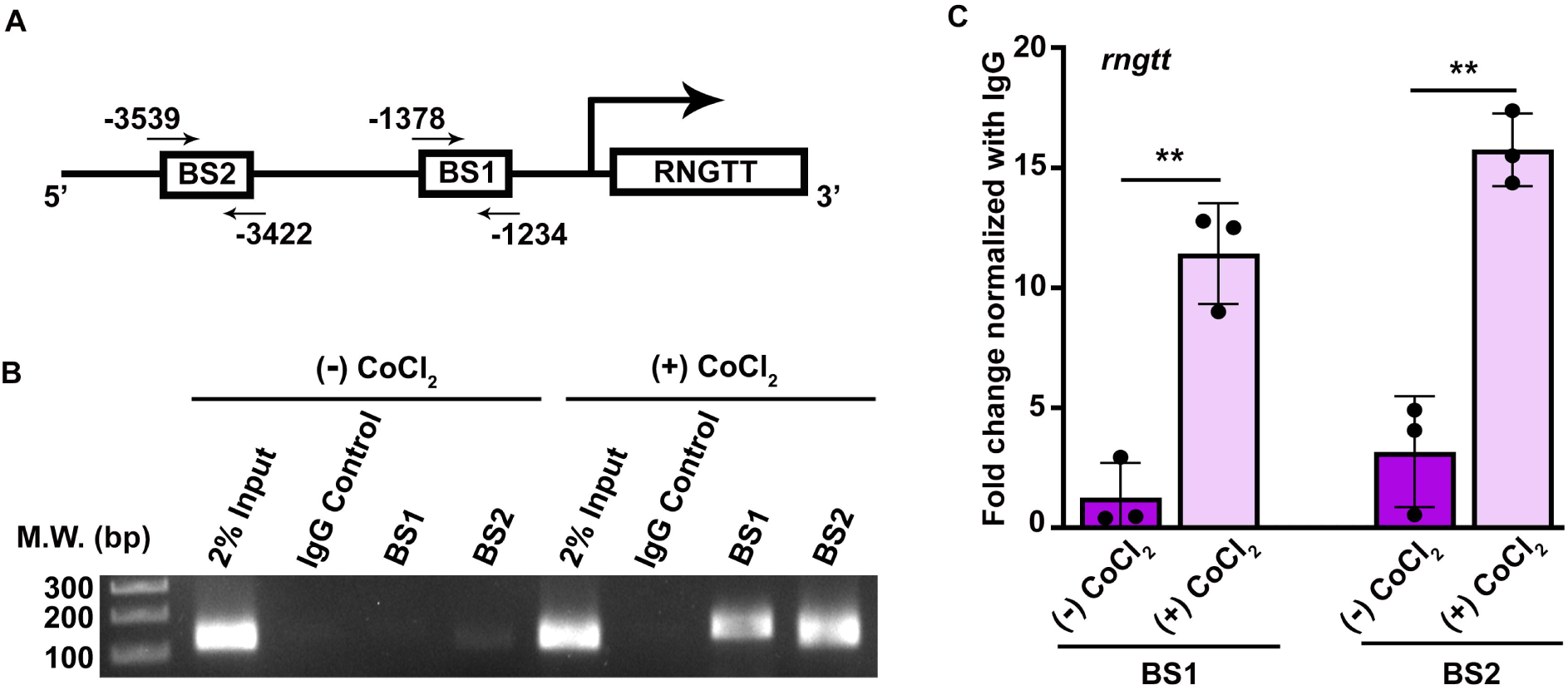
The promotor of mammalian CE (RNGTT) harbors hypoxia-responsive elements (HRE). A. Human RNGTT gene contains two HIF1α binding motif (R/CGTG) in its promoter region as shown in the schematic. B. Chromatin immunoprecipitation (ChIP) analysis showed direct binding of HIF1α in the *RNGTT* promoter regions (BS1 and BS2). 2% input samples were loaded. The molecular weight of the DNA ladder is shown on the left. C. ChIP assay followed by quantitative RT–PCR (RT–qPCR) showed ∼12 and 15-fold enrichment of HIF1α binding to the BS1 and BS2 of RNGTT promoter region in CoCl_2_ treated cells respectively. Statistical significance was calculated by unpaired two-tailed Student’s t-test represented as ± SD from three biological replicates. ns: non-significant, *P< 0.05, **P< 0.005, ***P< 0.0005, ****P< 0.0001; n≥3.

### 3.4 Depletion of HIF1**α** reduces expression of CE in osteosarcoma cells

To corroborate the ChIP assay results which showed binding of HIF1α to *RNGTT* promoter, HIF1α was depleted using pharmacological inhibitor PX478. PX478 is a well-known inhibitor of HIF1α (29). It affects its expression both at transcriptional, and translational levels. Moreover, PX478 prevents de-ubiquitination of ubiquitin-tagged HIF1α molecules (42). Osteosarcoma cells were treated with CoCl_2_ and PX478 was added to these cells only. To test if the binding of HIF1α to *RNGTT* promoter was direct or not, we performed ChIP assay as before (3.3) with PX478 treated or untreated CoCl_2_ induced hypoxic cells. The binding of HIF1α to BS1 and BS2 sites were markedly reduced in PX478 treated cells compared to untreated cells (Fig: S2A). Later, qPCR analysis showed that PX478 treatment led to ∼7 and 13-fold reduced binding of HIF1α to BS1 and BS2 sites respectively on *RNGTT* promoter as compared to PX478 untreated CoCl_2_ induced hypoxic cells. (Fig: S2B) thus confirming the direct binding of HIF1α to the *RNGTT* promoter.

Based on this result, we speculated the compromised binding of HIF1α to *RNGTT* promoter would result reduced expression of CE in PX478 treated cells. To test this, we treated U2OS cells and MG63 cells with CoCl_2_ or kept untreated. PX478 was added to CoCl_2_ induced hypoxic cells and total RNA and protein levels were measured from different treatment groups. We observed a reduction in the level of *RNGTT* mRNA in HIF1α inhibited, CoCl_2_ induced hypoxic samples compared to HIF1α non-inhibited CoCl_2_ induced hypoxic samples for both cell lines (Fig. 4A). Western blot analysis showed increased expression of HIF1α in CoCl_2_ induced hypoxic samples but decreased expression of HIF1α in PX478 treated, CoCl_2_ induced hypoxic samples in osteosarcoma cells (Figs. 4B-4E).

**Figure 4:**
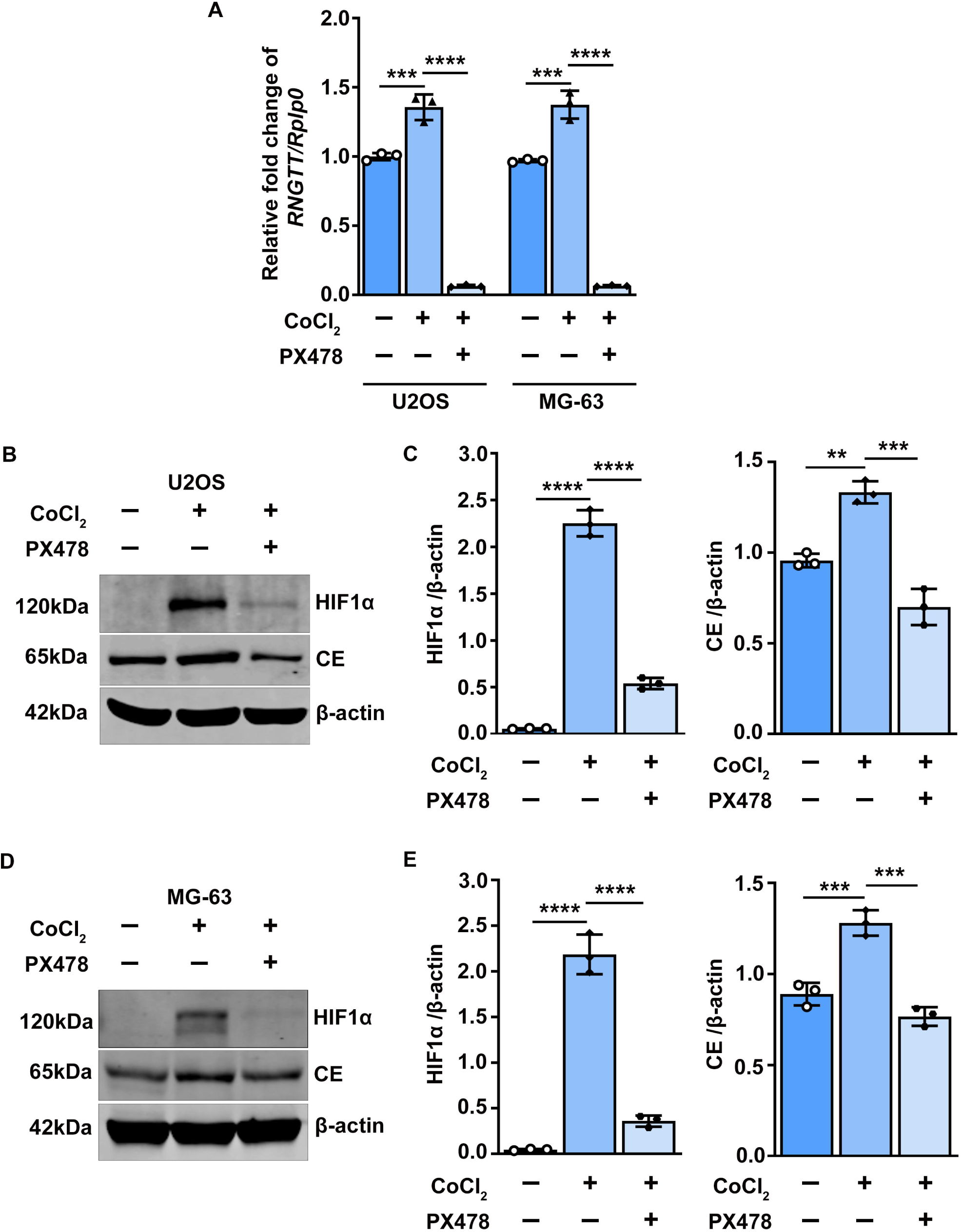
Depletion of HIF1α reduces expression of CE in osteosarcoma cells. A. Quantitative RT– PCR (RT–qPCR) analysis revealed treatment of osteosarcoma cells (U2OS and MG63) with 40µM HIF1α inhibitor PX478 in CoCl_2_ treated conditions resulting in a significant reduction in *RNGTT* mRNA expression. B & E. Western blot analysis shows an increased expressions of both HIF1α and CE protein upon CoCl_2_ treatment which were decreased significantly upon treatment with PX478 in presence of CoCl_2_. C & E. Quantification of HIF1α and CE protein by ImageJ software using three independent biological replicates. β-actin was used as the loading control. Values are represented as ± SD from three biological replicates. Statistical significance was drawn by performing unpaired two-tailed Student’s t-test, ns, P: non-significant, *P< 0.05, **P< 0.005, ***P< 0.0005, ****P< 0.0001; n≥3.

Next, we investigated the effect of HIF1α inhibition on the expression of CE. We observed CE expression also reduced in HIF1α inhibited samples as compared to respective controls in osteosarcoma cells (Figs. 4B-4E). Therefore, our study showed the elevated expression of *RNGTT* mRNA or the encoded protein during CoCl_2_ induced hypoxia. This was due to the direct binding of the HIF1α transcription factor on the *RNGTT* promoter which was abolished upon inhibition of HIF1α.

### 3.5 cCE stabilizes its targeted lncRNAs post-transcriptionally during hypoxia

The hypoxia microenvironment is one of the common features of solid tumors that regulates tumor cells both genotypically and phenotypically (43). Hypoxia, in turn, modulates the expression of various non-coding RNAs to make phenotypic changes in tumor cells, and lncRNAs are one of them (44,45). Previous findings from our group suggested a few lncRNAs could be targeted post-transcriptionally by cCE (21). Considering this fact, we are intrigued to understand whether cCE has any role in regulating its target lncRNAs during chemical induced hypoxia. For that, we have chosen ASH1L-AS1, GAS5-215, ENTPD3-AS1, NFYC-AS1, and H19-009 which are related to any solid tumors and a few of them established as substrates of cCE (21). H19-009 and GAS5 lncRNAs are elevated in various tumors (46,47). The other candidate lncRNAs are also involved in developing cancer occurrence and development; such as participation of ASH1L-AS1 in gastric cancer (GC), ENTPD3-AS1 in lung adenocarcinoma (LUAD), and renal cell carcinoma (RCC), NFYC-AS1 in LUAD (48–51).

To analyze the effect of hypoxia on these cCE-targeted lncRNAs, CoCl_2_ treated or untreated U2OS cells were separated into nuclear and cytoplasmic fractions efficiently as judged by the detection of Lamin A/C in the nuclear fractions and GAPDH in the cytoplasmic fractions (Fig. S3A). RNA was extracted from cytoplasmic fractions and relative expressions of the candidate lncRNAs were measured using RT-qPCR. We found that out of five lncRNAs, four of them (ASH1L-AS1, GAS5-215, ENTPD3-AS1, and H19-009) were significantly upregulated, whereas the expression of NFYC-AS1 was significantly reduced in CoCl_2_ treated cells (Fig: 5A). As the stability of some of these lncRNAs was regulated by cCE (20), we wondered if the upregulated expressions of these HRLs were contributed by cCE.

To examine this, we made Tetracycline-inducible stable U2OS cell lines expressing cytoplasmic restricted, catalytically inactive dominant negative form CE (referred to here as K294A), as schematically represented in Fig S3B. In this cell line, the nuclear localization signal (NLS) was deleted and the nuclear export signal (NES) was added to the CE construct to make it cytoplasm-restricted, as evident from the immunofluorescence image (Fig. 5B). These cells were treated with Doxycycline (Dox) to overexpress K294A (Fig: S3D). The same cell line was used as Control for subsequent assays where no overexpression was observed in absence of Dox (Fig. S3D). Control and K294A cells were treated with or without CoCl_2_ for 24 hrs as described earlier. Following incubation, all the cells were fractionated into nuclear and cytoplasmic fractions. The efficiency of fractions was assessed by western hybridization using antibodies against Lamin A/C and GAPDH as done earlier (Fig. 5C). Our data showed no cross-contamination among the fractions. The expression of CA9, was also checked to confirm the CoCl_2_ induced hypoxic condition (Fig. 5C). The steady-state expressions of the candidate lncRNAs in cytoplasmic fractions isolated from Control and K294A cells in normal condition, as well as CoCl_2_ induced hypoxic conditions, were measured using RT-qPCR (Fig. 5D).

**Figure 5:**
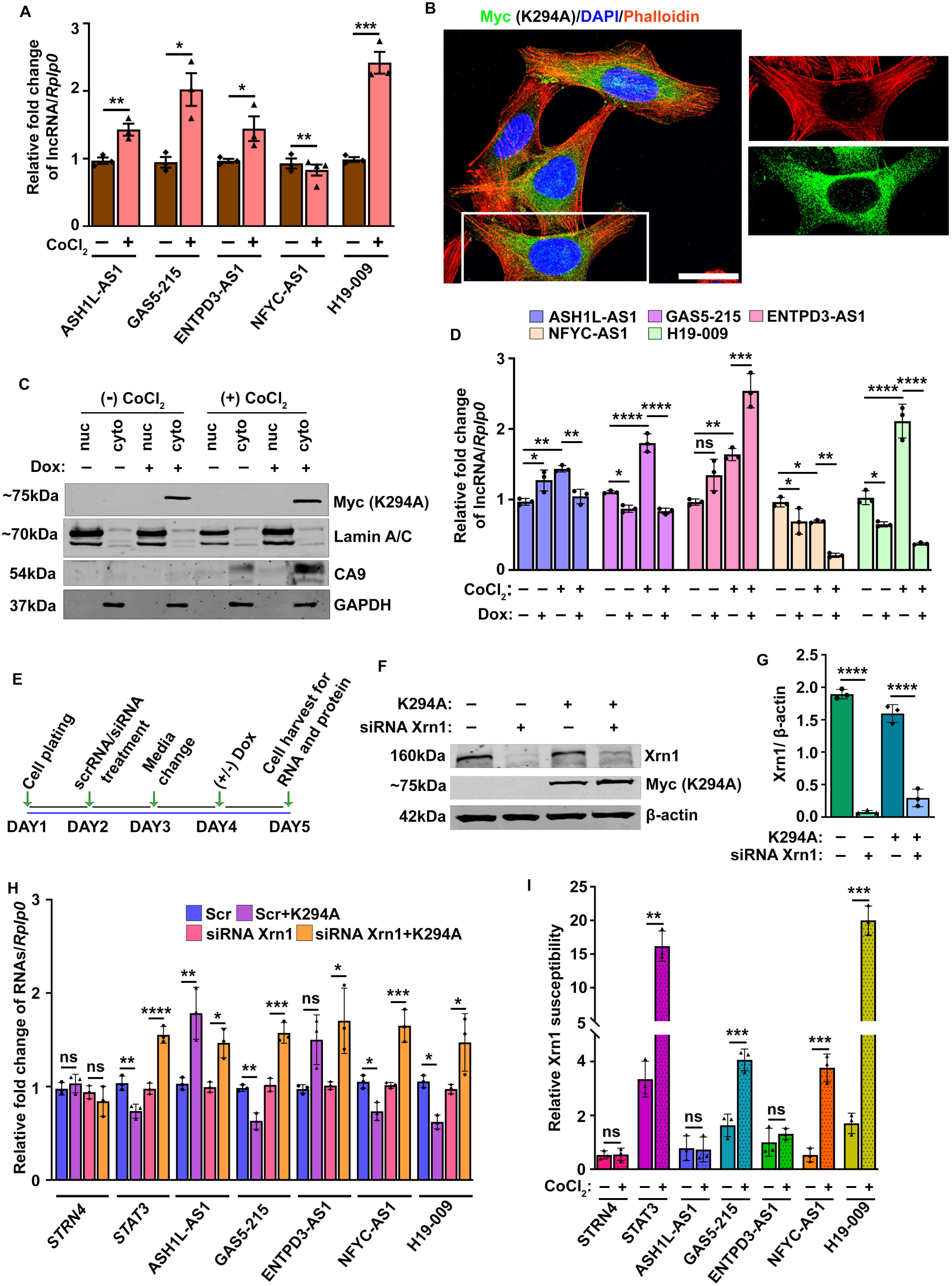
cCE stabilizes its targeted lncRNAs posttranscriptionally during hypoxia. A. RT-qPCR analysis of cCE-targeted lncRNAs shows differential expression patterns in cytoplasmic extracts of CoCl_2_ treated U2OS cells. The C_t_ values of each lncRNA was normalized to the internal control, *Rplp0*. The relative fold change of the transcripts was plotted after normalizing to the control, untreated samples. CoCl_2_ treatment significantly elevates the expression of ASH1L-AS1, GAS5-215, ENTPD3-AS1, and H19-009, whereas NFYC-AS1 expression was significantly reduced. A paired Two-tailed Student’s t-test was performed for statistical analysis. B. Confocal image of Doxycycline inducible stable clonal cell line C1 which expressed Myc tagged K294A in the cytoplasm only. C. Nuclear and cytoplasmic fractionation of U2OS cells expressing K294A in the presence or absence of CoCl_2_. Western blot of Myc and CA9 confirm K294A expression and CoCl_2-_induced hypoxia, respectively. The quality of fractionation was further substantiated by western blotting of nuclear marker (Lamin A/C) and cytoplasmic marker (GAPDH). D. The steady-state expressions of the candidate transcripts in K294A stable cells in ± CoCl_2_ conditions. GAS5-215, NFYC-AS1, and H19-009 were significantly decreased in K294A-expressing CoCl_2_ untreated cells. But ASH1L-AS1 was significantly increased and ENTPD3-AS1 did not show any significant change. Only GAS5-215 and H19-009 showed significantly reduced expression in CoCl_2-_treated K294A-expressing cells. The expression of each transcript was normalized to *Rplp0* and the relative fold change was plotted after comparing to control cells. E. Schematic of Xrn1 knockdown experiment. F. Western blot analysis of Xrn1 using cell lysate of K294A expressing/non-expressing cells transfected with either scramble or Xrn1 siRNA. G. Densitometric quantification of protein bands. β-actin was used as a loading control. H. RT-qPCR analysis of candidate transcripts using cytoplasmic RNAs of Xrn1 knockdown samples. *STRN4* is a control transcript whereas *STAT3*is a recapping target transcript respectively. The C_t_ values of each transcript was calculated by normalizing to the C_t_ values of the internal control, *Rplp0*. The relative fold change was calculated by normalizing to the control cells. Similar to STAT3, the steady-state levels of GAS5-215, NFYC-AS1, and H19-009 showed significant enrichment upon Xrn1 knockdown. Statistical significance was calculated using One way ANOVA to compare scr vs scr+K294A and si RNA Xrn1 vs si RNA Xrn1+ K294A. I. Xrn1 susceptibility assay was performed using poly (A) selected cytoplasmic RNAs from different conditions as described before (5D) and treated with ± Xrn1 followed by RNA extraction and RT-qPCR using gene-specific primers. Renila luciferase spike in control that was unchanged during ± Xrn1 was used as an internal control for normalization. The Xrn1 susceptibility is measured by differences in normalized C_t_ values between Xrn1 treated and non-treated sample. The Xrn1 susceptibility of cells expressing K294A and the non-expressing cells were plotted according to the previous study (Mukherjee et al., 2012). Unpaired Student’s t-test was used for statistical analysis. Values are represented as ± SD from three biological replicates. ns, P: non-significant, *P< 0.05, **P< 0.005, ***P< 0.0005, ****P< 0.0001; n≥3.

In agreement with our previous study (21), no significant changes were observed in the expression of ENTPD3-AS1 whereas ASH1L-AS1 showed significant elevation in its expression in K294A-expressing cells (Fig. 5D). However, in contrast to our earlier observation with transiently transfected cells (21), we noticed significantly reduced expressions of GAS5-215, NFYC-AS1, and H19-009 in K294A cells as compared to Control cells under similar conditions (Fig. 5D). This difference may be attributed to the usage of clonal stable cell lines in the current study expressing the similar level of K294A in contrast to a mixed population of cells expressing different levels of K294A along with untransfected cells generated from transient transfection experiments performed in the previous study (21). A similar trend was also observed for mRNA substrates of cCE, as reported in previous studies where decreased expressions of cCE targeted mRNAs were noticed in U2OS cell line stably expressing K294A (18,52). We hypothesized since K294A cells express inactive cCE, the cCE-targeted transcripts like GAS5-215, NFYC-AS1, and H19-009 present in the cytoplasm could not be recapped and would be susceptible to exoribonuclease-mediated degradation resulting in reduced expression in K294A cells. To address this, we depleted Xrn1, the major exoribonuclease from K294A and Control cells using specific siRNA targeting Xrn1 as shown in the schematic Fig: 5E. Cytoplasmic fractions were isolated to collect RNA and protein from these treatment groups. Our data showed more than 80% depletion of Xrn1 (Figs: 5F-G). Further qPCR analysis of cytoplasmic RNAs from the above treatment groups revealed elevated expressions of GAS5-215, NFYC-AS1 and H19-009 in K294A cells compared to Control cells when Xrn1 was depleted (Fig. 5H) supporting our hypothesis. On the other hand, ASH1L-AS1 and ENTPD3-AS1 might be protected by some other mechanism that yet to be studied.

Interestingly, we observed the expressions of ASH1L-AS1, GAS5-215 and H19-009, which were significantly upregulated in CoCl_2_ induced hypoxic control cells, were reduced in CoCl_2_ induced hypoxic K294A cells (Fig. 5D). In addition, the expression of NFYC-AS1 that did not elevate in response to CoCl2 treatment, was reduced markedly in presence of K294A (Fig. 5D). The possible reason could be during hypoxic stress, these transcripts might have lost its 5’ cap structure and in the absence of active cCE, they could not be recapped, subjected to exoribonucleases. Hence, their expressions were reduced in CoCl_2_ mimetic hypoxic K294A cells (Fig. 5D). To examine this possibility, we performed Xrn1 susceptibility assay that can distinguish uncapped RNA from capped RNAs (16). Cytoplasmic poly(A) RNAs from CoCl_2_ treated and untreated cells mixed with capped Rennila luciferase mRNA were subjected to *in vitro* Xrn1 digestion followed by RT-qPCR for 5 candidate lncRNAs, *STRN4* and *STAT3*, which were negative and positive controls of cCE respectively. The C_t_ values obtained for each transcript in ± CoCl_2_ treated cells were normalized against that of the spike luciferase mRNA, which remained unchanged by Xrn1 treatment.

Xrn1 susceptibility or loss of 5’ ends are represented as the difference in relative Ct values before and after Xrn1 digestion of the same transcript in ±K294A cells as described in the methods (Fig: 5H). We observed in absence of active cCE, GAS5-215, H19-009 and NFYC-AS1 transcripts lost their caps during CoCl_2_ mimetic hypoxic conditions as evidenced by high susceptibility to Xrn1 similar to STAT3 transcript suggesting role of cCE protecting these transcripts during CoCl_2_ treatment. In contrast, expression of ASH1L-AS1 transcript that was reduced in CoCl_2_ treated K294A expressing cells, remained unaffected by Xrn1 digestion, indicating retaining cap structure under same condition (Fig: 5H). The similar notion was observed for expression of ENTPD3-AS1 transcript too that increased in K294A expressing cells, retained its cap structures under the same condition like STRN4 (Fig: 5H). These data suggest that cCE plays an important role in regulating the stability of GAS5-215, H19-009 and NFYC-AS1 transcripts in CoCl_2_ mimetic hypoxic conditions post-transcriptionally.

## 4. Conclusion

Nuclear capping is a co-transcriptional event whereas cytoplasmic capping is a post-transcriptional event. In our previous study, we demonstrated that a subset of lncRNAs is post-transcriptionally stabilized by cCE which recapped the uncapped transcripts in the cytoplasm. In this study, we showed for the first time that CE/cCE is upregulated by HIF1α due to the binding of this transcription factor to HRE-containing sites in the *RNGTT* promoter. We also examined some tumor-associated lncRNAs and found they are differentially expressed during CoCl_2_ induced hypoxia. Importantly, a few of them (H19-009, GAS5-215, and NFYC-AS1) are post-transcriptionally regulated by cCE as evidenced by their decreased expressions in K294A cells under normal conditions. Additionally, we observed reduced expressions of these transcripts in CoCl_2_ induced hypoxic K294A cells which suggests cCE post-transcriptionally regulates the stability of these transcripts. Further, we showed cCE protects it’s target transcripts, H19-009, GAS5-215, and NFYC-AS1 during hypoxia when they lost their caps. However, no significant change in Xrn1 susceptibility was seen for ASH1L-AS1 under this condition. Perhaps, other mechanisms may be responsible for its down regulation which needs to be studied in the future.

In this present study, we have examined the effect of hypoxia on the total pool of CE using CoCl_2_ as a hypoxia mimetic agent on osteosarcoma cells and hTERT RPE-1 as cellular models. Mechanistically, the enhanced expression of CE/cCE under this chemical induced hypoxia is executed by HIF1α which is a known transcription factor and the master regulator of hypoxia. CE is known to maintain the stability of most RNA pol II transcripts that include both mRNAs and lncRNAs. Further investigation should be carried out to identify all hypoxia-responsive RNAs regulated by cCE. Since the present study relies upon only on CoCl_2_ that stimulated HIF1α-mediated responses via its inhibition of prolyl hydroxylase (39), and this treatment might have some off target effects. It would be better for future experiments to be conducted under physical hypoxic condition that mimics more closely the tumour microenvironment to examine the effect of hypoxic stress on CE/cCE and the plausible post transcriptional regulation of the transcriptome by CE/cCE.

Moreover, earlier study showed the protective function of cCE on its target transcripts during recovery from a brief arsenite induced oxidative stress through recapping the decapped transcripts. Further studies should examine if cCE play similar protective role to its target transcripts during recovery from hypoxic conditions. Taken together, our data reveals HIF1α mediated upregulation of CE/cCE and a novel function of cCE in post-transcriptional regulation of target transcripts during CoCl_2_ induced hypoxia.

## Acknowledgments

This work was supported by extramural grants CRG/2019/006427 and CRG/2023/006384 from the Department of Science and Technology, Government of India to Dr. Chandrama Mukherjee. The authors also thank UGC, Government of India for a UGC-SRF fellowship to Mr. Safirul Islam and the Department of Biotechnology, India for a Ramlingaswami Re-entry fellowship to Dr. Chandrama Mukherjee. The authors thank the Institute of Health Sciences, Presidency University for the departmental instrument facility and Presidency University for necessary infrastructural support, Dr. Piyali Mukherjee, Presidency University for providing Lamin antibody, Dr. Shubhra Majumder, Presidency University for providing access to Zeiss Microscope and Dr. Abhik Saha, Presidency University for providing access to Licor imager.

## Author contributions: CRediT

**Safirul Islam:** Conceptualization, Methodology, Validation, Investigation, Formal Analysis, Visualization, Writing-original draft, Writing - Review & Editing; **Chandrama Mukherjee:** Conceptualization, Resources, Writing - Original Draft, Writing - Review & Editing, Supervision, Project administration, Funding acquisition.

## List of Abbreviations

CE: mRNA capping enzyme
cCE: cytoplasmic mRNA capping enzyme
HIFs: hypoxia inducible factors
HRR: hypoxia responsive RNA
CoCl_2_: Cobalt chloride
ChIP: chromatin immunoprecipitation
LncRNA: long noncoding RNA
miRNA: microRNA
circRNA: circular RNA
RNGTT: RNA Guanylyltransferase And 5’-Phosphatase
RNMT: RNA guanine 7-methyl transferase
RAMAC: RNA guanine 7-methyl transferase activating subunit
K294A: dominant negative mutant of cCE
bHLH: basic-helix–loop–helix
LUAD: lung adenocarcinoma
RCC: renal cell carcinoma

