## Supplementary Information for "Hypoxia inducible factor HIF1α elevates expression of mRNA capping enzyme during cobalt chloride-induced hypoxia"

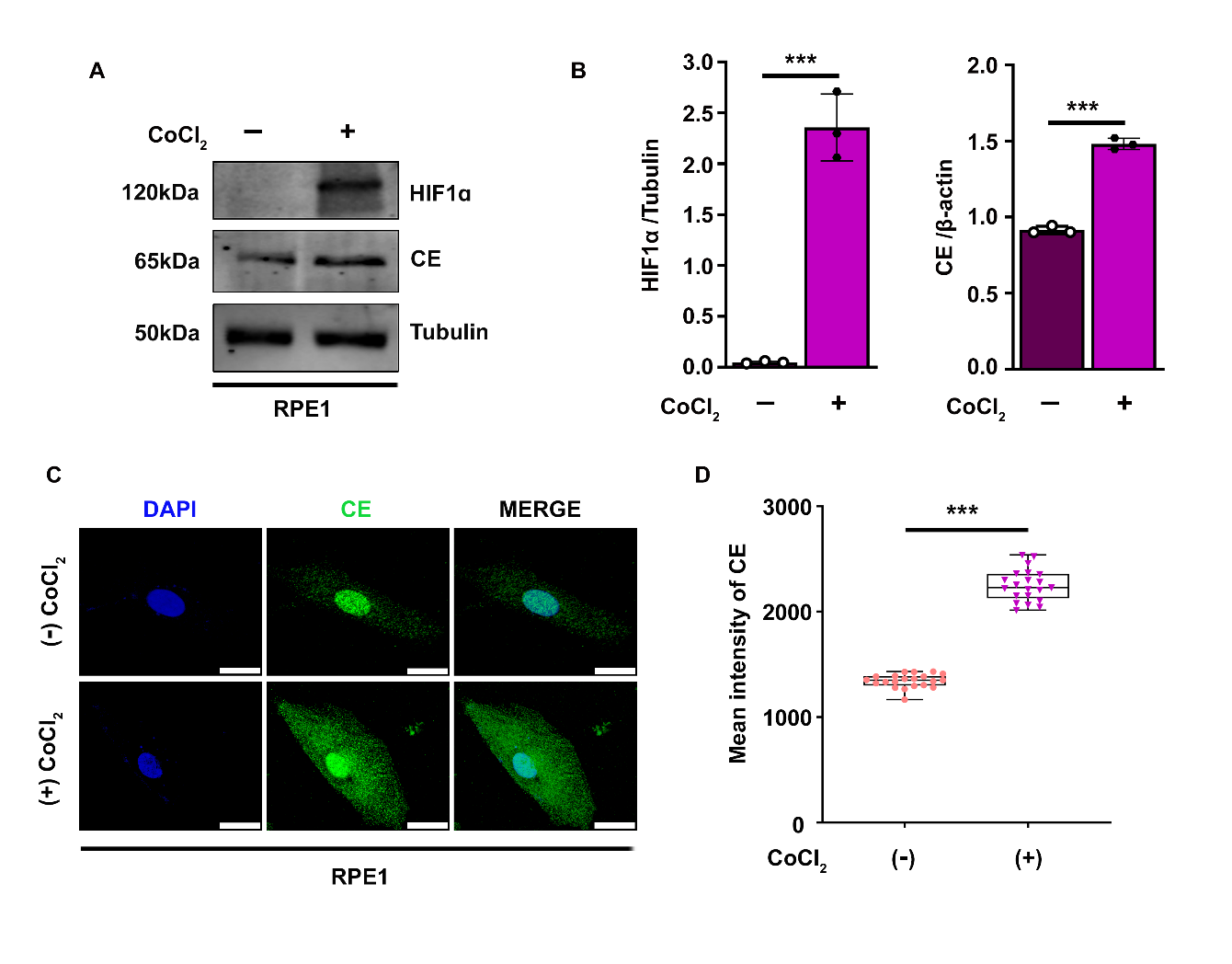
**Supplemental Figure 1: Cobalt chloride-induced hypoxia elevates mRNA-capping enzyme (CE) expression in RPE1 cells.** A. Western blot analysis shows a significant increase in the expression of CE in CoCl_2_ induced hypoxic RPE1 cell lysate. HIF1α blot confirms CoCl2 induced hypoxia induction. Tubulin was used th loading control. B. Quantification of HIF1α and CE protein by ImageJ software using three independent biological replicates. Plot shows the fold change in the expression of CE compared Tubulin. C. Immunostaining analysis showing enhanced expression of CE in CoCl_2_ induced hypoxic RPE1 cells as compared to untreated cells. The scale bar represents 50µm. D. Quantification of mean CE intensity using Zeiss Zen 3.3 software package, n≥20 cells. Values are represented as ± SD from three biological replicates using unpaired two-tail Student’s *t*-tests statistical analysis. ns, P: non-significant, *P< 0.05, **P< 0.005, ***P< 0.0005, ****P< 0.0001; n≥3.

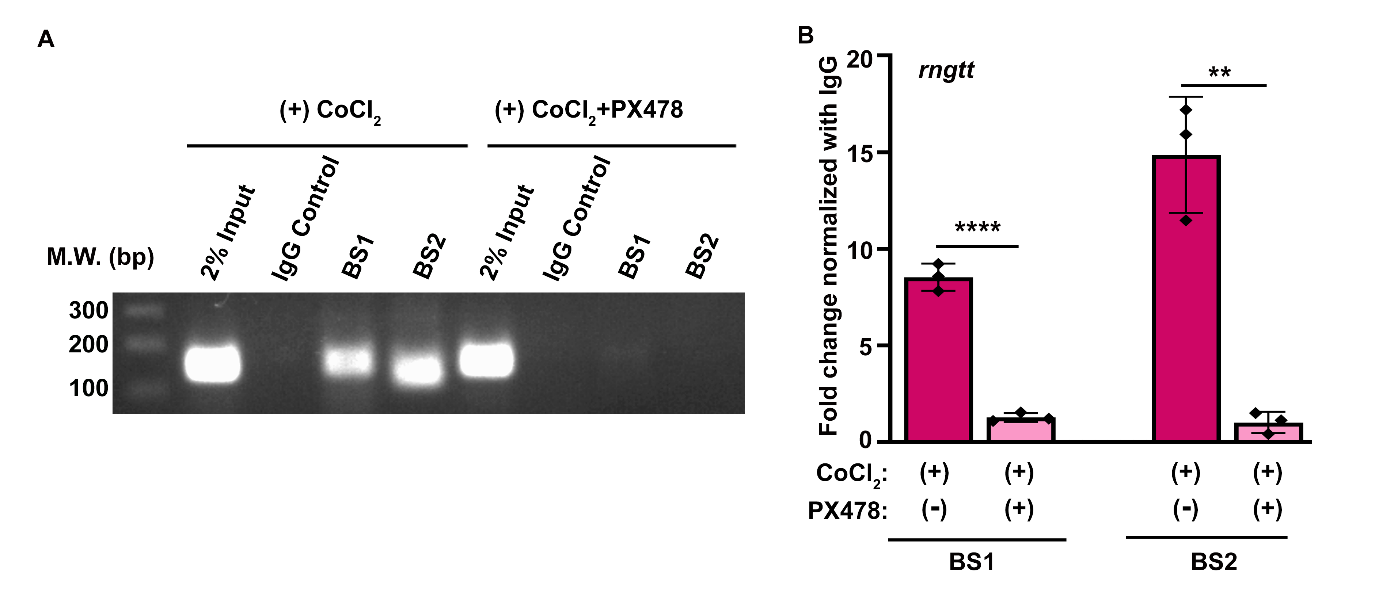

**Supplemental Figure 2: HIF1α fails to bind to mammalian CE (*RNGTT*) promoter region (BS1 and BS2) in presence of PX478.**

A. Chromatin immunoprecipitation (ChIP) analysis showed loss of HIF1α binding to *RNGTT* promoter region (BS1 and BS2) in presence of HIF1α pharmacological inhibitor PX478. 2% input samples were loaded. The molecular weight of the DNA ladder is shown on the left. B. ChIP assay followed by quantitative RT–PCR (RT–qPCR) showed ∼7 and 13-fold decrease of HIF1α binding to the BS1 and BS2 of *RNGTT* promoter region in CoCl_2_ +PX478 treated cells respectively as compared to CoCl_2_ treated cells. Values are represented as ± SD from three biological replicates using unpaired two-tail Student’s *t*-tests statistical analysis. ns: non-significant, showing the fold change normalized with IgG, *P< 0.05, **P< 0.005, ***P< 0.0005, ****P< 0.0001; n≥3.

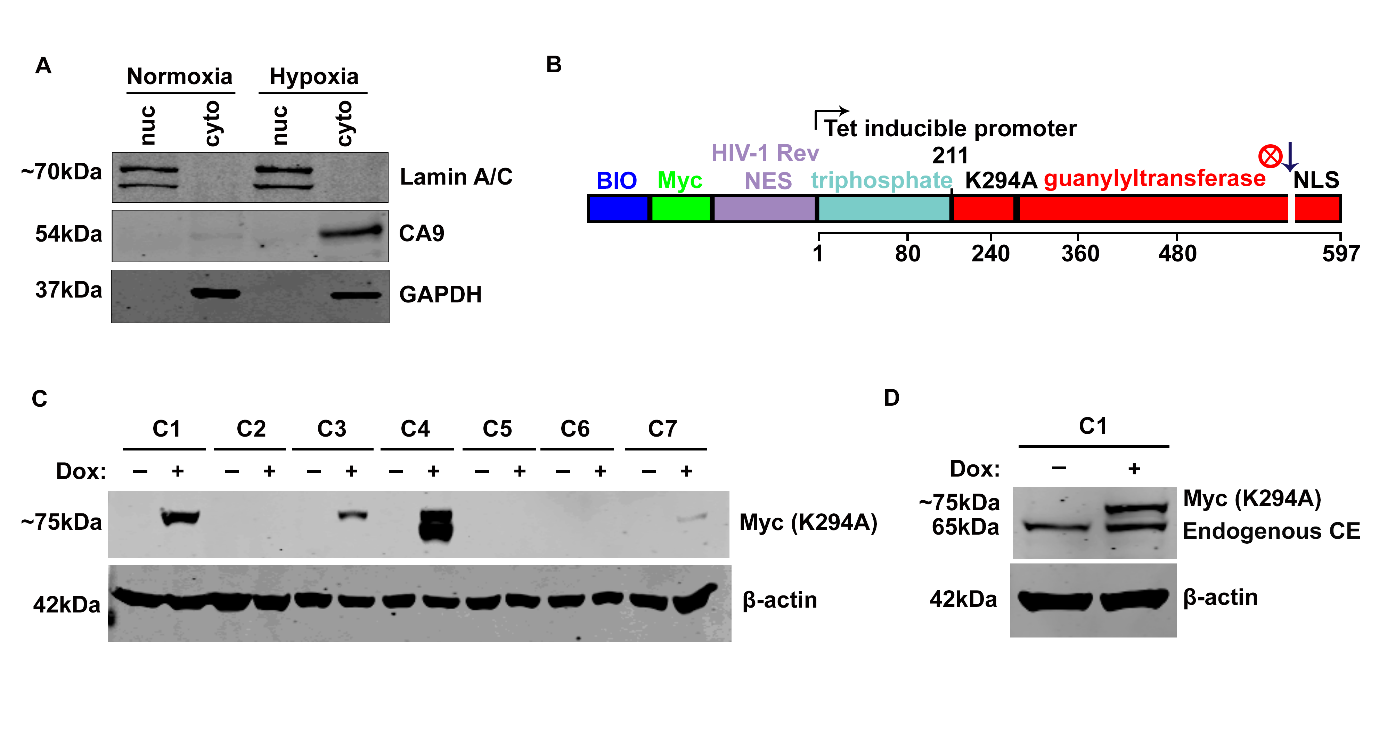

**Supplemental Figure 3: Generation of Tet- inducible stable cell line expressing K294A**

A. Biochemical fractionation of ± CoCl_2_ treated U2OS cells into nuclear/ cytoplasmic fractions. The quality of fractionation was assessed by immunoblotting of nuclear marker (Lamin A/C) and cytoplasmic marker (GAPDH). CA9 immunoblotting was done to confirm the CoCl_2-_induced hypoxic condition. B. Schematic representation of the dominant negative form of cytoplasmic restricted capping inhibited construct (K294A) Bio and Myc denote the biotinylation and Myc tag respectively. The nuclear export sequence (NES) of HIV is inserted in the N-terminus and the Nuclear Localization Signal (NLS) from the C-terminus is deleted to restrict expression of CE only in the cytoplasm. The lysine (K) residue at 294 positions is mutated to alanine (A) to make the construct catalytically inactive. C. Western blot of Myc (K294A) using cytoplasmic lysate from different clones (C1-C7) of K294A stable cells in presence of Doxycycline. β-actin was used as a loading control. D. Western blot of Myc (K294A) and endogenous CE using lysate from C1, K294A stable cells using anti-CE antibody that shows tight regulation of K294A expression only in presence of Doxycycline.

**Table S1: List of primers**

| **Primer Name** | | | | **Primer Sequence (5’ 3’)** | **Accession number** |
| --- | --- | --- | --- | --- | --- |
| **mRNA primer sequences (Homo sapiens)** | | | | | |
| HIF1α | | Forward | | TGACCCTGCACTCAATCAAG | NM_001530.4 |
|  |  | Reverse | | GGCTCAGGTGAACTTTGTCTAG |  |
| CA9 | | Forward | | ATCAGAAGAAGAGGGCTCCC | NM_013083.2 |
|  |  | Reverse | | TCCATAGCGCCAATGACTCT |  |
| RNGTT | | Forward | | TCATCAATTCTGTGGCTGGG | NM_003800.5 |
|  |  | Reverse | | CCGAGTACCATCTGCTTTCCA |  |
| Rplp0 | | Forward | | GGAGAAACTGCTGCCTCATATC | NM_053275.4 |
|  |  | Reverse | | CAGCAGCTGGCACCTTATT |  |
| STAT3 | | Forward | | TAGCAGGATGGCCCAATGGAATCA | NM_001369512.1 |
|  |  | Reverse | | AGCTGTCACTGTAGAGCTGATGGA |  |
| STRN4 | | Forward | | AACCTGGCCGGCAATCAGAGAGGAT | NM_001039877.2 |
|  |  | Reverse | | CCCAAACACCACTCTCCTGGACAATTA |  |
| GFP spike-in | | Forward | | TTTCACTGGAGTTGTCCCAAT |  |
|  |  | Reverse | | TTCCGTATGTTGCATCACCT |  |
| Rluc spike-in | | Forward | | GGATGATAACTGGTCCGCAG |  |
|  |  | Reverse | | GGCCGCGTTACCATGTAAAA |  |
| Uncapped GFP RNA (spike in control) | | UUU CAC UGG AGU UGU CCC AAU UCU UGU UGA AUU AGA UGG CGA UGU UAA UGG GCA AAA AUU CUC UGU CAG UGG AGA GGG UGA AGG UGA UGC AAC AUA CGG AA | | | |
| **LncRNA Primer Sequence (Homo sapiens)** | | | | | |
| **Primer Name** | | | | **Primer Sequence (5’ 3’)** | **Transcript ID** |
| H19-009 | | Forward | | CGTTCCAGGCAGAAAGAGCAA | ENST00000422826.1 |
|  |  | Reverse | | GGACCCCTCTGTCCTGTGT |  |
| GAS5-215 | | Forward | | GCACTATCAGTTCACTGCAACC | ENST00000442067.5 |
|  |  | Reverse | | CAGCCTGGACAACATAGCTGTA |  |
| NFYC-AS1 | | Forward | | CAGGCAAGAGCACAACATGG | ENST00000606277.1 |
|  |  | Reverse | | AGTTTCTGTGCGAGGGTGAT |  |
| ENTPD3-AS1 | | Forward | | CGTCTGCTACACCCAACTGA | ENST00000420850.1 |
|  |  | Reverse | | TCCTTGAGTAGAGGCCAAGC |  |
| ASH1L-AS1 | | Forward | | CGGTTGACCTGAGCCTACTT | ENST00000456633.1 |
|  |  | Reverse | | AGCGGTACAATACCCGTGTC |  |
| **Primers used in ChIP assay** | | | | | |
| **Primer Name** | | | **Primer Sequence (5’ 3’)** | | |
| BS1 | Forward | | TGAGCTACTGGGTATAGCGTG | | |
|  | Reverse | | AAGTGGTAAAAACCTTCCATCCC | | |
| BS2 | Forward | | ACTACAGGCACGTCACCACA | | |
|  | Reverse | | CCAAGGCAGGCAGATCATTTG | | |
| **Primers used in Cloning** | | | | | |
| K294A cloning primer | Forward | | CCTCGTAAAGAATTCATGGATCAATGCAGAAGCTG | | |
|  | Reverse | | AGGTGGGGATCCTCAGGTTGGCCGATGCAGTCT | | |
